# Hydrodynamics shapes self-recruitment in anemonefishes

**DOI:** 10.1101/2022.06.11.495394

**Authors:** Masaaki Sato, Kentaro Honda, Yohei Nakamura, Lawrence Patrick C. Bernardo, Klenthon O. Bolisay, Takahiro Yamamoto, Eugene C. Herrera, Yuichi Nakajima, Chunlan Lian, Wilfredo H. Uy, Miguel D. Fortes, Kazuo Nadaoka, Masahiro Nakaoka

## Abstract

Many marine species have a pelagic larval phase that undergo dispersal among habitats. Studies on marine larval dispersal have revealed a large variation in the spatial scale of dispersal, and accumulated evidence has shown that seascape patchiness is the major determinant for variation in self-recruitment. However, few studies have investigated the influence of geographic settings on marine larval dispersal. Bays or lagoons generally enhance the retention of larvae, while larvae are more likely to be flushed by strong currents in open coasts. To examine associations between larval dispersal, geographic setting, and hydrodynamics, we compared fin-scale dispersal patterns, self-recruitment, and local retention of two anemonefishes (*Amphiprion frenatus* and *A. perideraion*) between a semi-enclosed bay and an open coast in the Philippines combining genetic parentage analysis and biophysical dispersal modelling. Contrary to our expectations, parentage analysis revealed lower estimates of self-recruitment in the semi-closed bay (0–2%) than in the open coast (14–15%). The result was consistent with dispersal simulations predicting lower local retention and self-recruitment in the former (0.4% and 19%) than in the latter (2.9% and 38%). Dispersal modelling also showed that cross-shore currents toward offshore were much stronger around the semi-closed bay and were negatively correlated with local retention and self-recruitment. These results suggest that stronger cross-shore currents around the semi-closed bay transport anemonefish larvae to the offshore and mainly contributed to the lower self-recruitment. Our results highlight difficulty in predicting self-recruitment from geographic setting alone and importance of hydrodynamics on it.

## Introduction

Many marine species have life cycles with a pelagic larval phase that undergoes dispersal among habitat patches, and a benthic adult phase that occurs in discrete habitat patches after larval settlement (Jones et al., 2009). Marine larvae are subjected to oceanographic processes that transport them at varying distances from their original birthplace (Cowen and Sponaugle, 2009). Additionally, their behavior affects the success of dispersal and settlement (Paris and Cowen, 2004). Quantifying the dispersal patterns of marine species is essential for predicting population dynamics and managing marine populations.

The controversy over whether marine populations are open or closed has continued among marine ecologists for a long time period (Mora and Sale, 2002). While marine ecologists traditionally believed that marine populations were open over hundreds to thousands of kilometers (Mora and Sale, 2002), empirical studies have provided a new paradigm in which marine populations are relatively closed and can be maintained by local replenishment (Jones et al., 1999; Swearer et al., 1999). After this paradigm shift, the application of genetic parentage analysis to marine systems has contributed significantly in clarifying dispersal patterns of coral reef fishes (Jones et al., 2005; Saenz-Agudelo et al., 2011; D’Aloia et al., 2013). Comparisons of marine dispersal estimates using this method revealed a large variation in the spatial scale of dispersal in coral reef fishes. For example, estimates of dispersal distance ranged from less than 50 m to 35 km for *Amphiprion spp*., and from 1 to 48 km for *Chaetodon vagabundus* (Jones et al., 2005; Berumen et al., 2012; Saenz-Agudelo et al., 2012; Abesamis et al., 2017). Marine dispersal studies have also shown a rapid decline in the probability functions of dispersal success within the first few kilometers (Buston et al., 2012; Saenz-Agudelo et al., 2012). Larval recruitment to the natal patch can be measured in two ways: self-recruitment and local retention (Nanninga et al., 2015). self-recruitment denotes the fraction of all incoming recruits to a specific patch produced at that patch, whereas local retention describes the fraction of all larvae produced at a focal patch returning to that patch (Fig. 1) (Botsford et al., 2009). Because of the inherent nature of what is measured using parentage analysis (i.e., the fraction of all recruits that have been produced by local parents), empirical studies typically report estimates of self-recruitment, ranging from 0 to 65% for coral reef fishes (Jones et al., 2005; Saenz-Agudelo et al., 2011; D’Aloia et al., 2013; Nanninga et al., 2015). Pinsky et al. (2012) suggested that seascape patchiness, the spatial configuration of habitat patches, is one of the major determinants of self-recruitment variation among populations. The study predicted that isolated populations in patchy seascapes have higher self-recruitment than populations in continuous seascapes because of the lack of larval supply from external populations. This prediction is largely consistent with the high self-recruitment in isolated islands (Almany et al. 2007; Planes et al. 2009; Berumen et al. 2012; also see, Nanninga et al. 2015).

**Fig. 1.**
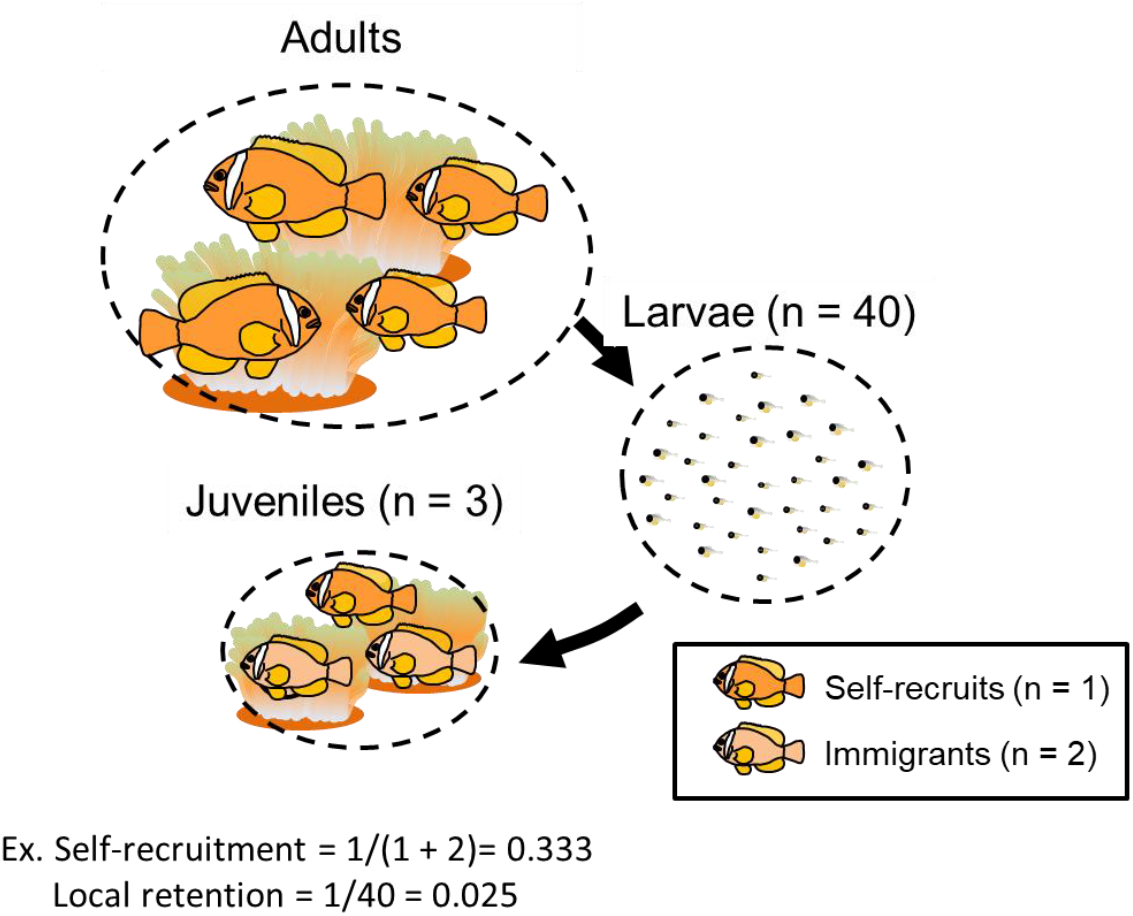
Schematic diagram shows self-recruitment and local retention in a population of *Amphiprion frenatus*. Self-recruitment is defined as the number of self-recruits divided by the total number of juveniles (self-recruits and immigrants). Local retention is defined as the number of self-recruits divided by the total larval rereleases from that population.

Although seascape patchiness is an important factor for predicting the self-recruitment of marine larvae, the geographic characteristics of coastal systems (i.e., open coast vs. bay) and associated water circulation can also influence larval transport and dispersal patterns in marine organisms. Bays or lagoons generally have restricted connections to the open sea and exhibit long residence times of water (Kjerfve and Magill, 1989; Sponaugle and Cowen, 2002), which may enhance the local retention and self-recruitment of larvae. Meanwhile, pelagic larvae are flushed by strong currents and can be easily dispersed or transported far away along open coasts (Swearer and Shima, 2010; Huebert et al., 2011). Therefore, the local retention and self-recruitment are expected to vary between the different geographic settings. However, there are limited examples of comparison of larval dispersal patterns between contrasting geographic settings (but also see, Saenz-Agudelo et al. 2011). Understanding the relationship between dispersal patterns and geographic setting is helpful for better management planning, such as size decisions and spatial setting of marine protected areas (MPAs).

Hydrodynamic models verified using oceanographic data have been used as a basis for larval dispersal simulations (Cowen et al., 2007; Werner et al., 2007; Cowen and Sponaugle, 2009). Recent dispersal models integrate biological characteristics of larvae such as larval mortality, competitive period, and migration behavior (Cowen et al., 2000, 2006; Nanninga et al., 2015; Treml et al., 2015). Such models are especially effective for exploring larval dispersal processes when combined with field/empirical methods (Werner et al., 2007). Meanwhile, most studies have used genetic differentiation measures as empirical estimates of larval dispersal for comparison with the physical predictions of dispersal (White et al., 2010; Nakajima et al., 2014; Hawkins et al., 2019). Such genetic differentiation measures indicate averaged population connectivity across generations over evolutionary timescales, and are normally useful for determining genetic structures over large spatial scales in the sea (hundreds to thousands of kilometers). In contrast, genetic parentage analysis enabled us to clarify fine-scale patterns of larval dispersal over ecological time scales, whereas there are still few integrations of this approach with hydrodynamic model (but also see, Nanninga et al., 2015). Comparing the modeling and empirical results of ecological measures of larval dispersal, both consistencies and mismatches between them provide insights into the mechanisms influencing larval dispersal as well as implications for management strategies.

In this study, we compared the dispersal patterns, self-recruitment, and local retention of two anemonefishes (*Amphiprion frenatus* and *A. perideraion*) between a semi-enclosed bay and an open coast located in continuous seascapes using genetic parentage analysis and biophysical modeling. Anemonefish have been widely used as model species for genetic parentage analysis, mainly because they are easily located and can be caught underwater through use of SCUBA as well as their relatively short PLD makes it easy to elucidate local dispersal patterns (e.g., Planes et al. 2009; Berumen et al. 2012; Sato et al. 2017). Sato et al. (2017) showed the dispersal patterns and self-recruitment of the two species on the open coast of the Philippines. The present study empirically estimated the dispersal patterns and self-recruitment in the semi-enclosed bay in the same country for comparison. Biophysical models were then developed to simulate larval dispersal and calculated local retention and self-recruitment of the two anemonefishes in the two sites. The following questions were addressed: (1) Is there variation in self-recruitment estimates of parentage analysis between a semi-enclosed bay and an open coast? (2) Does the larval dispersal simulation using a biophysical model confirm empirical estimates of parentage analysis? (3) How is hydrodynamics related to larval dispersal patterns?

## Materials

### Study species and study sites

Our target species were the tomato anemonefish (*Amphiprion frenatus*) and the pink anemonefish (*A. perideraion*), which occur throughout the eastern Indian Ocean to the western Pacific Ocean. The two study sites are situated on fringing reefs around Puerto Galera (13°30’N, 120°57’E) in the Verde Island Passage and around Laguindingan (8°38’N, 124°28’E) in the southern Bohol Sea, Philippines (Fig. 2) (Sato et al., 2014a, 2017). Puerto Galera has a semi-enclosed bay, which connects to the passage via two channels, that is, the Manila Channel in the northwest and the Batangas Channel in the northeast, while Laguindingan has an open coast facing the Bohol Sea. The study area of both sites is part of a marine protected area (MPA) that is strongly protected against fishing activity (Sato et al., 2014a, 2017).

**Fig. 2.**
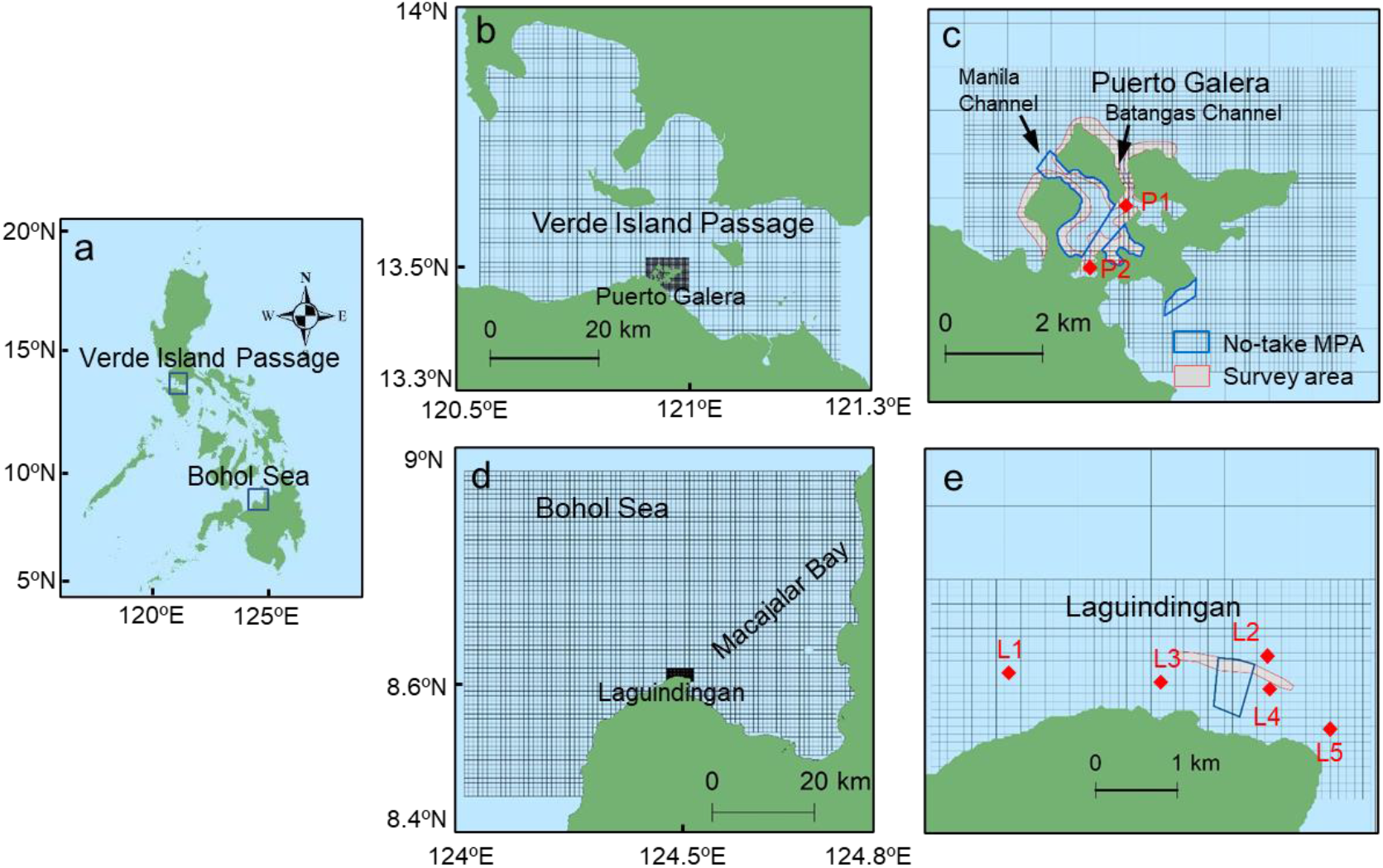
**a** Map of the Philippines. **b**–**c** Modeled domains of the Verde Island Passage and semi-enclosed bay of Puerto Galera (meshed grids) with sensor array locations (red diamond: P1 and P2). **d**–**e** Modeled domains of the Bohol Sea and open coast of Laguindingan (meshed grids) with sensor array locations (red diamond: L1–L5). The outer and inner domains of both models have a cell size of 900 m and 100 m, respectively. The locations of MPAs, survey areas, prominent embayment, and channels are also indicated.

### Field collection of genetic samples in Puerto Galera

A previous study estimated the larval dispersal and self-recruitment of the target anemonefishes in the open coast of LD (Sato et al. 2017), whereas the present study investigated their dispersal patterns in the semi-closed bay of Puerto Galera for comparison. Using the same procedure employed by Sato et al. (2017), we collected genetic samples of target anemonefish around Puerto Galera (Fig. 3). In June 2012, we conducted a preliminary survey by snorkeling on coral reefs in Puerto Galera to record the location of anemonefish and host anemones, because they were abundant only in such habitats (Sato et al., 2014). A GPS device (Garmin eTrex 30) was used to determine the locations. On the basis of the location data, we captured anemonefish using hand-nets and clove oil and measured their TL to the nearest mm underwater using SCUBA in September 2012. Anemonefish were fin clipped using scissors and then released back into the same host sea anemone. Fish that were too small to be fin clipped (<30 mm) were collected. We sampled a total of 206 *A. frenatus* and 98 *A. perideraion* at Puerto Galera in September 2012. Based on the classification of Sato et al. (2017), a total of 48 and 17 juveniles (<30 mm), 24 (30–46 mm) and 12 non-breeders (30–39 mm), and 134 (>46 mm) and 69 breeders (>39 mm) were collected from 86 and 51 host anemones for *A. frenatus* and *A. perideraion*, respectively. This practice resulted in 97.1 % and 95.8 % of breeders collected within the study area for the respective species. A previous study estimated that 30 mm *Amphiprion* are approximately three to four months old, respectively (Ochi, 1986). Therefore, the collected juveniles (<30 mm) were predicted to disperse from May to September mainly during the Southwest monsoon.

**Fig. 3.**
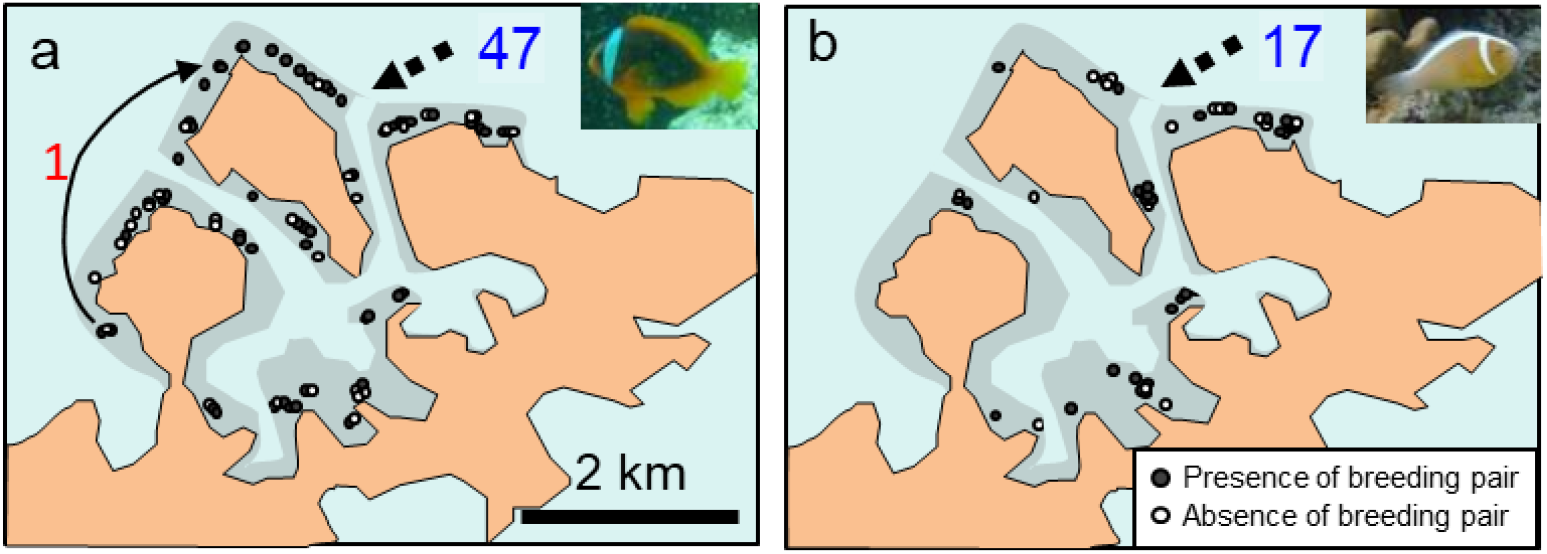
Maps for sample collection of anemonefishes around a semi-closed bay of Puerto Galera, northern Mindoro Island, the Philippines. **a** *Amphiprion frenatus* (n = 206) and **b** *A. perideraion* (n = 98) in the study area (gray area: 5-15m). Dispersal tracks of self-recruits and its individual number are shown by an arrow and red number, while immigrants from the outside and its individual number are shown by dashed arrows and blue numbers.

To estimate the pelagic larval duration (PLD) of each species, lapilli otoliths were dissected from the subsamples of juveniles. The PLDs of each fish was determined by counting the number of daily increments of the otolith using an otolith measurement system (Ratoc System Engineering Inc.; www.ratoc.com). PLDs were estimated to be 7–14 days for *A. frenatus* and 8–15 days for *A. perideraion* (n = 5 for each).

### Parentage analysis for Puerto Galera populations

We extracted genomic DNA, amplified fragments, and sequenced and scored them according to Sato et al. (2014b). The individuals were genotyped using 22 microsatellite loci for *A. frenatus* and 24 loci for *A. perideraion* (Table S1, see Supplemental Information). We conducted a genetic parentage analysis for Puerto Galera samples to identify self-recruits of each target species using the program COLONY v. 2.0.6.4 (Jones and Wang, 2010). This program implements a full-likelihood parentage analysis method and defines the priori probability that the true parent is present in the samples. COLONY is robust to uncertainty in the sampling rate of true parentage and has been shown to outperform other programs (Harrison et al. 2013). We tested a range of sampling rates of the true parent (0.05–0.30) and used 0.05 as a conservative sampling proportion for both species (D’Aloia et al., 2013). Other settings for the analysis in COLONY were as follows: full-likelihood method, medium run length, medium probability precision, no inbreeding, and assumed monogamy for both sexes. We informed our COLONY runs with allele frequencies estimated from the sampled individuals. To assess the information sufficiency of our markers for the accurate reconstruction of parental assignment, we also used the simulation module in COLONY (see Supplemental Information).

Parentage analyses were run to test the pool of juveniles (n = 48 for *A. frenatus* and n = 17 for *A. perideraion*) against candidate mothers (n = 68 for *A. frenatus* and n = 28 for *A. perideraion*) and fathers (n = 66 for *A. frenatus* and n = 41 for *A. perideraion*), with a mistyping rate of 1 % to account for genotyping errors. We only accepted parentage assignments with a probability of > 0.8.

Once the assignment was made, we regarded assigned juveniles as self-recruits in the study site, and remaining juveniles as immigrants from outside of the study site, calculating the self-recruitment at Puerto Galera as follows:

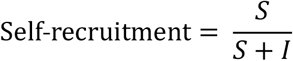

where *S* was the number of settlers assigned to breeders in the study site (self-recruits) and *I* was the number of settlers not assigned to the breeders (immigrants) (Fig. 1).

### Hydrodynamic model

Using Delft3D-Flow, a 3D-capable hydrodynamic simulation program (Deltares, 2014a), we developed two hydrodynamic models, i.e., the Puerto Galera model extending from 13.38 to 13.93 N and from 120.62 to 121.26 E, and the Laguindingan model extending from 8.41 to 8.99 N and from 124.07 to 124.78 E (Fig. 2). Both models were comprised of two computational domains using one-way nesting technique to properly incorporate the effects of the complicated topographic setup and tidal fluctuation. For the Puerto Galera model, the Verde Island Passage area was an outer domain with a cell size of 900 m and the Puerto Galera domain with a cell size of 100 m was the inner domain. For the Laguindingan model, the Bohol Sea area was an outer domain with a cell size of 900 m, and the Laguindingan area with a cell size of 100 m was the inner domain. Thirteen and fifteen vertical Z-layers were used for Puerto Galera and Laguindingan models, respectively. The General Bathymetric Chart of the Oceans (GEBCO: http://www.gebco.net/) data were supplemented with the past bathymetry data and echo sounder measurements in both areas, where coral reefs make topography highly variable. Open water boundaries for both models were forced with tidal variation derived from the PH model (Pokavanich 2009) and with salinity and water temperature from JCOPE2 (Miyazawa et al., 2009) and Hybrid Coordinate Modeling System analysis data (HYCOM GLBu0.08) (https://hycom.org/dataserver/glb-analysis). The water surface was forced with a spatially uniform wind, ocean heat exchange, and precipitation.

The Puerto Galera model simulation was run with the forcing conditions from February 1 to September 30, 2012, with a spin-up period of one month while the Laguindingan model was simulated from January 1 to July 31, 2013, with the same spin-up period. The time step was set to 6 s for both models. Details of the hydrodynamic models and evaluation of model performance were provided in Supplemental Information (Figs. S1, S2, and Table S2).

### Larval dispersal simulation

We performed larval dispersal simulations for anemonefish juveniles (<30 mm) at the two sites using Delft3D-PART (Deltares, 2014b). PART computed the position of every individual particle by advection and diffusion, and simulated transport processes by means of a particle-tracking method using flow data from the FLOW module. The processes were assumed to be deterministic except for a random displacement of the particle at each time step. Time step for the particle-tracking calculation was set to 30 minutes. PART simulated larval dispersal from the study site and external surrounding sources into the focal area of the study site in each model domain. The main model output of interest was local retention (the fraction of all released particles from the study site that settled back to there) and self-recruitment (the proportion of settling particles that are derived from study site) for each study site (Fig. 3).

Based on the spawning biology of *Amphiprion* and the expected hatching period for the collected juvenile (<30 mm) in both sites, we started the particle releases on the day of spring tide four months prior to the sample collection (9 spring tides for Puerto Galera and 11 spring tides for Laguindingan). Based on the distance at which 90 % of anemonefish larvae recruit (17.6 km) as estimated by Catalano et al. (2021), we utilized the 20 km radius to set source sites around each study site. In the Puerto Galera model, a total of 75,600 particles were released from 84 locations (100 particles × 9 spring tides × 84 locations) along the shore within a radius of 20 km from the anemonefish survey area in Puerto Galera (Fig. 4). Meanwhile, in the Laguindingan model, we released 33,000 particles from 30 locations (100 particles × 11 spring tides × 30 locations) along the shore within a radius of 20 km from the survey area of Laguindingan (Fig. 5). Biological input values of anemonefishes were derived from the literature, and the measurement of target species or closely related species (see, Supplemental Information). Pelagic larval duration was set at 15 days for target anemonefishes based on our otolith analysis.

**Fig. 4.**
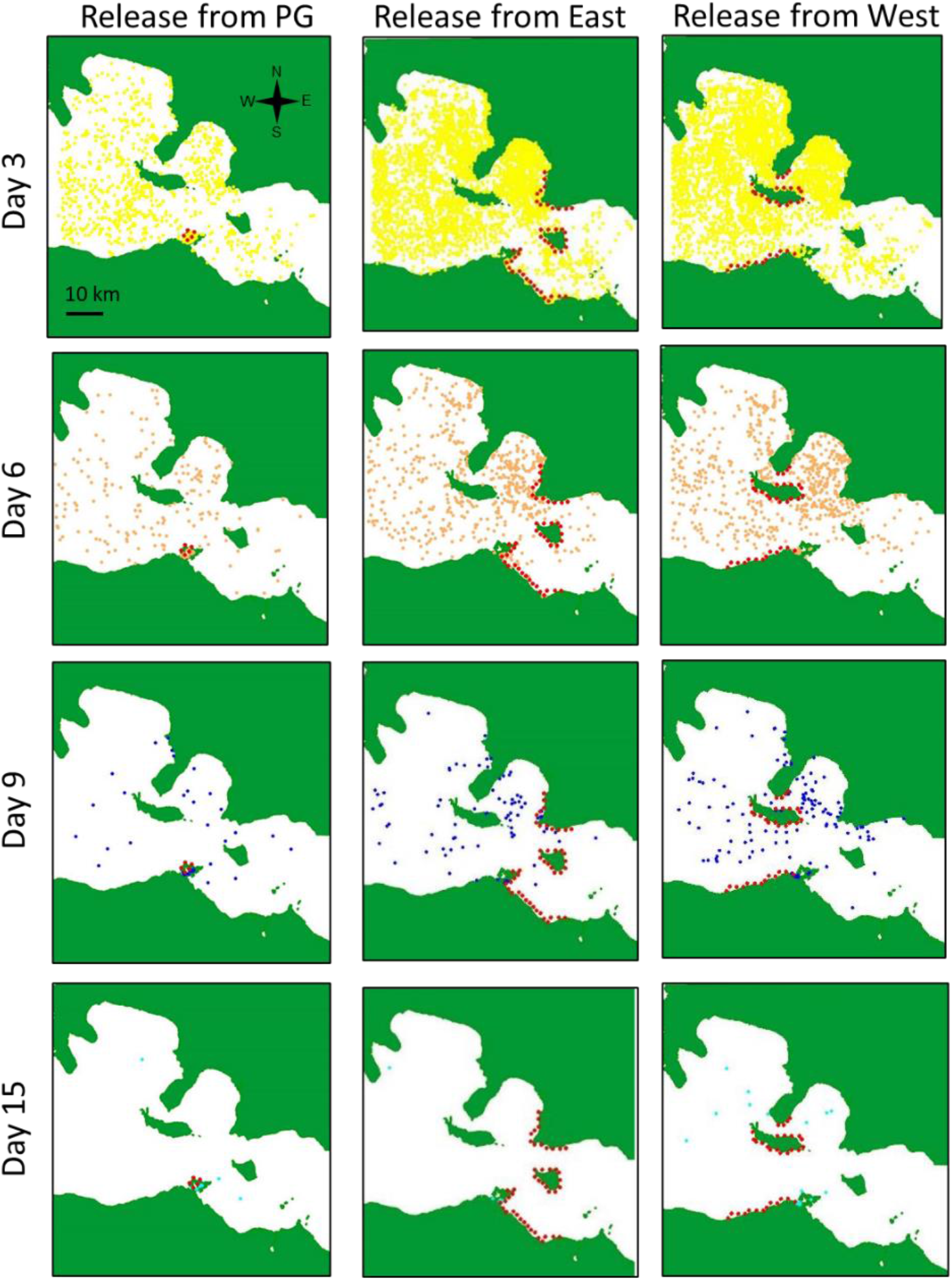
Distribution of particles released from Puerto Galera (PG), the eastern and western areas of potential habitats within a radius of 20km from the focal study area, predicted by biophysical dispersal simulation. We showed distribution of particles in the entire water column 3, 6, 9, 15 days after releases on May 6, May 21, June 4, June 20, July 4, July 19, August 2, August 18, and August 31, 2012. Anemonefish larvae are predicted to settle on sedentary habitats during competent period from Day 6 to Day 15. Red circles indicate release locations.

**Fig. 5.**
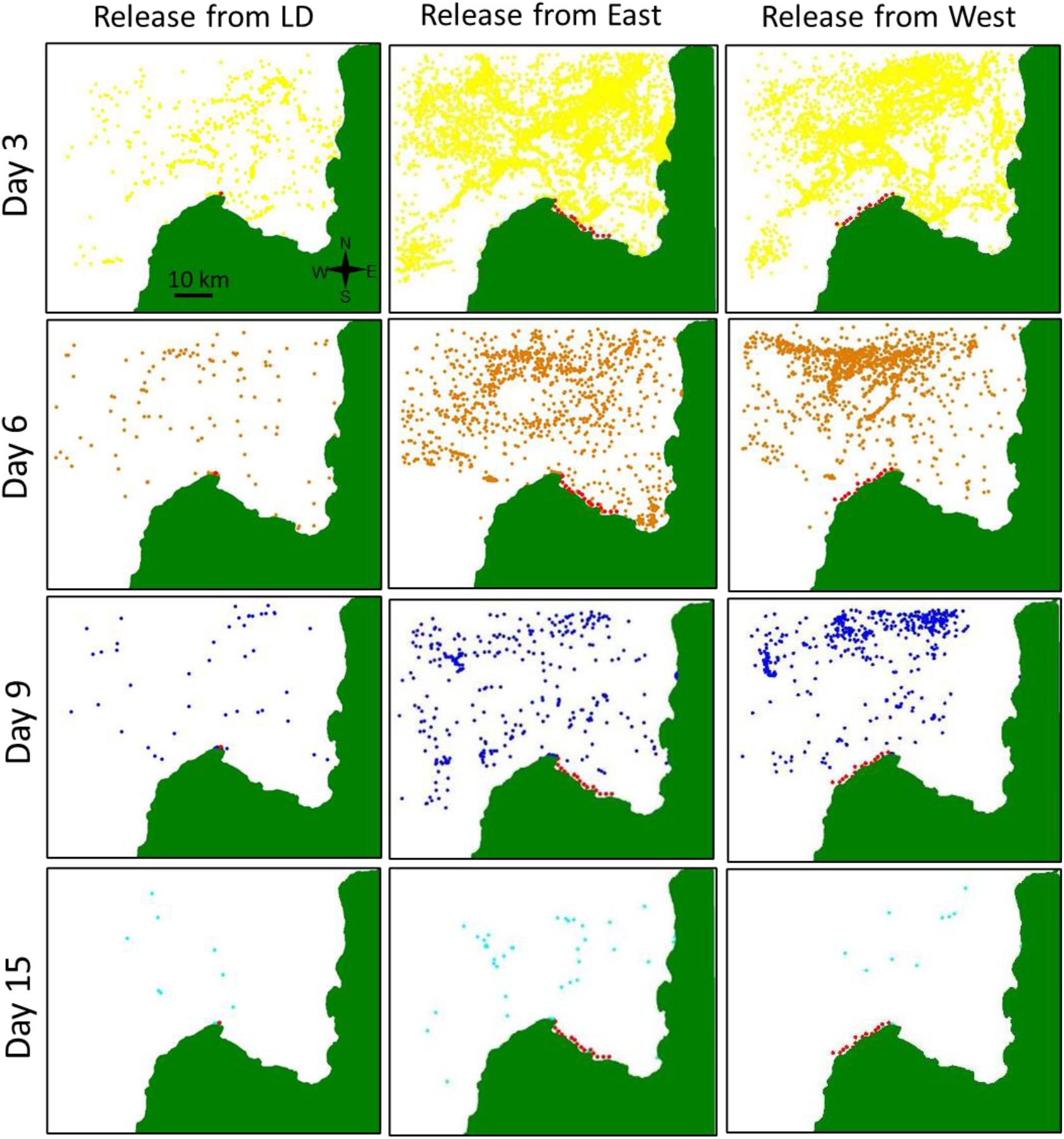
Distribution of particles released from Laguindingan (LD), the eastern and western areas of potential habitats within a radius of 20km from the study area, predicted by biophysical dispersal simulation. We showed distribution of particles in the entire water column 3, 6, 9, 15 days after releases on February 10, February 26, March 12, March 27, April 10, April 26, May 10, May 25, June 9, Jun23, July 8, 2013. Anemonefish larvae are predicted to settle on sedentary habitats during competent period from Day 6 to Day 15. Red circles indicate release locations.

Because biological traits of larvae affect dispersal, we assessed the model’s sensitivity to different biological parameter inputs according to Nanninga et al. (2015). Five parameters were examined for their effect on local retention and self-recruitment: competitive period, horizontal sensory zone, larval mortality, number of particle releases from the study site, and vertical migration behavior. Three competitive periods were set for larvae, where larvae were allowed to start settling after 6 days (±2) (Dixson et al., 2011). Horizontal sensory navigation was simulated by creating retention zones of 1 (+1/–0.5) km radius around the settling areas. Larval mortality was calculated with a half-life of 6 (±2) days. The number of particle releases from the study site per spring tide were set at 100 particle releases (+100 particles/–50 particles) per source location. We adopted two vertical migration behaviors (passive and empirical). The empirical migration behavior includes diel and ontogenetic vertical migration (see, Supplemental information), which was based on distribution patterns measured via plankton surveys in the same family (Irisson et al., 2010; Huebert et al., 2011). Dispersal simulation in the original scenario (competency = 6day, sensory zone = 1km, morality = a half-life of 6 days, vertical migration = empirical migration, number of particles = 100 particles) was repeated for three times to check the variation of the local retention and self-recruitment.

### Influences of hydrodynamics on local retention and self-recruitment

To examine influences of hydrodynamics on local retention and self-recruitment, a multiple regression analysis was conducted using the simulation results of Delft 3D-flow and Delft3D-PART. We calculated 6 days-averaged residual velocity of vertically averaged flow at all grids in the inner domains of the Puerto Galera and Laguindingan models (Figs. 2, S3, and S4). The residual velocity was obtained by averaging the flow velocity over 6 subsequent days from each date of particle releases (n = 9 for Puerto Galera and n = 11 for Laguindingan). Then, those values were averaged over the all grids, and the North-South (N-S) and East-West (E-W) components were utilized for this analysis. We assessed the local retention and self-recruitment in relation to the mean N-S and E-W velocities using a generalized linear model. The local retention and self-recruitment for each date of particle release in the original simulation scenario of Delft3D-PART were utilized as dependent variables (9 spring tides × 3 replicates for Puerto Galera and 11 spring tides × 3 replicates for Laguindingan). For the analysis, a mixture of binomial and gamma distributions, which is called a gamma hurdle model,” was used for both local retention and self-recruitment as in previous studies (Sato et al. 2021) because these values are continuous and contain zero values. We used BIC weight to average each model parameter found within the model set with ΔBIC < 4 (Burnham and Anderson, 2004). This model averaging framework was adapted to estimate robust parameters (Johnson and Omland, 2004). We also obtained the relative importance of each variable by summing the BIC weights over the model set with ΔBIC < 4.

## Results

### Parentage analysis in Puerto Galera

The parentage analysis of COLONY revealed that only one *A. frenatus* juvenile was assigned to a breeder within the study area of Puerto Galera (Fig. 3). This result indicates that the percentages of self-recruitment in Puerto Galera was only 2.1 % (one self-recruit/48 juveniles) for *A. frenatus* juveniles, while it was 0 % (zero/17 juveniles) for *A. perideraion* juveniles.

### Larval dispersal model

Larval dispersal simulation showed that local retention and self-recruitment were markedly higher in Laguindingan than in Puerto Galera. In the original simulation scenario (n = 3), the levels of local retention were 0.4 ± 0.1 % (mean ± SD) in the Puerto Galera model and 2.9 ± 0.2 % in the Laguindingan model, and those of self-recruitment were 19.0 ± 2.7 % and 37.9 ± 1.7 %, respectively. The Puerto Galera model predicted that only a small number of the particles from Puerto Galera were retained (18.0/4500), while a larger number of particles from the western side settled to Puerto Galera (71.9/30600), mainly contributing to a lower self-recruitment there (19.0 %) (Fig. 4 and Table S3). The Laguindingan model predicted that there were a large number of particles from Laguindingan locally settled there (32.3/1100) compared to those from the east (48.6/16500) and west (4.1/15400), resulting in a higher self-recruitment (37.9 %) in Laguindingan (Fig. 5 and Table S3).

Individual parameter perturbation (IPP) of biological traits had variable effects on simulated levels of local retention in Puerto Galera and Laguindingan (Table 1). Changes to settlement competency, sensory zone, or mortality rate led to variations in local retention. Maximum scenarios yielded almost five-and two-times higher estimates of local retention (2.1 % and 5.3 %) than the original simulation scenarios (0.4 % and 2.9 %) in Puerto Galera and Laguindingan models, respectively; while minimum scenarios yielded half and one fourth (0.2 % and 0.8 %) of the estimates in respective models. Predicted self-recruitment in both models also varied through changes in the number of self-recruits and imports from surrounding sources according to different competitive periods, sensory zones, number of particle releases from the study site, and vertical migration behavior (Table 1). Maximum scenarios yielded about 2.5 and 1.5-times estimates (46.5 % and 57.0 %) of the original values of self-recruitment (19.0 % and 37.9 %) in Puerto Galera and Laguindingan models, respectively; while minimum scenarios yielded approximately an half and three fifth (8.9 % and 21.6 %) of the original estimates in respective models.

**Table 1.** Effects of individual parameter perturbation (IPP) on local retention (LR) and self-recruitment (SR) at Puerto Galera and Laguindingan.

|  | Original value | IPP range | Negative IPP (%) |  |  | Positive IPP (%) |  |  |
| --- | --- | --- | --- | --- | --- | --- | --- | --- |
|  |  |  |  | LR | SR |  | LR | SR |
| Puerto Galera |  |  |  |  |  |  |  |  |
| Competency | 6 day | ±2day | 4 d : | 2.1 | 33.5 | 8 d : | 0.2 | 17.9 |
| Sensory zones | 1km | +1 km/-0.5 km | 0.5 km : | 0.3 | 32.4 | 2 km : | 0.5 | 19.6 |
| Mortality | 6 day | ±2day | 4 day : | 0.3 | 20.4 | 8 day : | 0.5 | 20.5 |
| Vertical migration | Empirical migration | Passive migration | Passive : | 0.4 | 46.5 |  |  |  |
| Number of particles* | 100 particles | +100 particles/-50 particles | 50 particles : | 0.3 | 8.9 | 200 particles : | 0.5 | 35.0 |
| Parameter combination | n/a | Max/min retention potential | Min: | 0.2 | 8.9 | Max: | 2.1 | 46.5 |
| Laguindingan |  |  |  |  |  |  |  |  |
| Competency | 6 day | ±2day | 4 d : | 5.3 | 26.7 | 8 d : | 1.1 | 30.4 |
| Sensory zones | 1km | +1 km/-0.5 km | 0.5 km : | 2.8 | 63.0 | 2 km : | 3.0 | 32.4 |
| Mortality | 6 day | ±2day | 4 day : | 2.1 | 37.9 | 8 day : | 3.5 | 37.7 |
| Vertical migration | Empirical migration | Passive migration | Passive : | 0.8 | 21.6 |  |  |  |
| Number of particles* | 100 particles | +100 particles/-50 particles | 50 particles : | 2.8 | 22.7 | 200 particles : | 3.2 | 57.0 |
| Parameter combination | n/a | Max/min retention potential | Min: | 0.8 | 21.6 | Max: | 5.3 | 57.0 |

To test model sensitivity, biological parameters of the Delft PART simulations were systematically perturbed for Puerto Galera and Laguindingan models. Original values of local retention at Puerto Galera and Laguindingan were $0.4 \pm 0.1$ % and $2.9 \pm 0.2$ %, respectively, while those of self-recruitment at respective site were $19.0 \pm 2.7$ % and $37.9 \pm 1.7$ %. \*We only perturbed number of particles releases from the focal areas in Puerto Galera and Laguindingan. Refer to text for parameter descriptions.

### Influences of hydrodynamics on local retention and self-recruitment

Multiple regression analysis indicated the continuous (gamma) portion of the mean N-S and E-W velocities have high relative importance (RI > 0.65) for local retention and self-recruitment (Table 2). The model averaging results showed that the local retention and self-recruitment were increased with increasing the mean E-W velocity, but these values were decreased with increasing the mean N-S velocity (Fig. 6).

**Table 2.** Model averaging results showing the averaged coefficients, standard errors (SE), and relative importance (RI) of parameters for the Gamma hurdle models.

| Variables | Coefficient | SE | RI |
| --- | --- | --- | --- |
| Local retention |  |  |  |
| Intercept<sup>a</sup> | -3.36 | 0.18 | - |
| E-W velocity<sup>a</sup> | 11.13 | 2.96 | 0.97 |
| N-S velocity<sup>a</sup> | -46.08 | 5.17 | 0.97 |
| zi(Intercept)<sup>b</sup> | -2.37 | 0.58 | - |
| zi(E-W velocity)<sup>b</sup> | 13.37 | 12.63 | 0.15 |
| zi(N-S velocity)<sup>b</sup> | -24.24 | 30.04 | 0.13 |
| Self-recruitment |  |  |  |
| Intercept<sup>a</sup> | -1.20 | 0.17 | - |
| E-W velocity<sup>a</sup> | 7.16 | 2.43 | 0.85 |
| N-S velocity<sup>a</sup> | -10.32 | 4.11 | 0.66 |
| zi(Intercept)<sup>b</sup> | -2.38 | 0.56 | - |
| zi(E-W velocity)<sup>b</sup> | 13.37 | 12.63 | 0.10 |
| zi(N-S velocity)<sup>b</sup> | -24.24 | 30.04 | 0.09 |

<sup>a</sup>Continuous (gamma) portion of parameters. <sup>b</sup>Hurdle (binomial) portion of parameters.

**Fig. 6.**
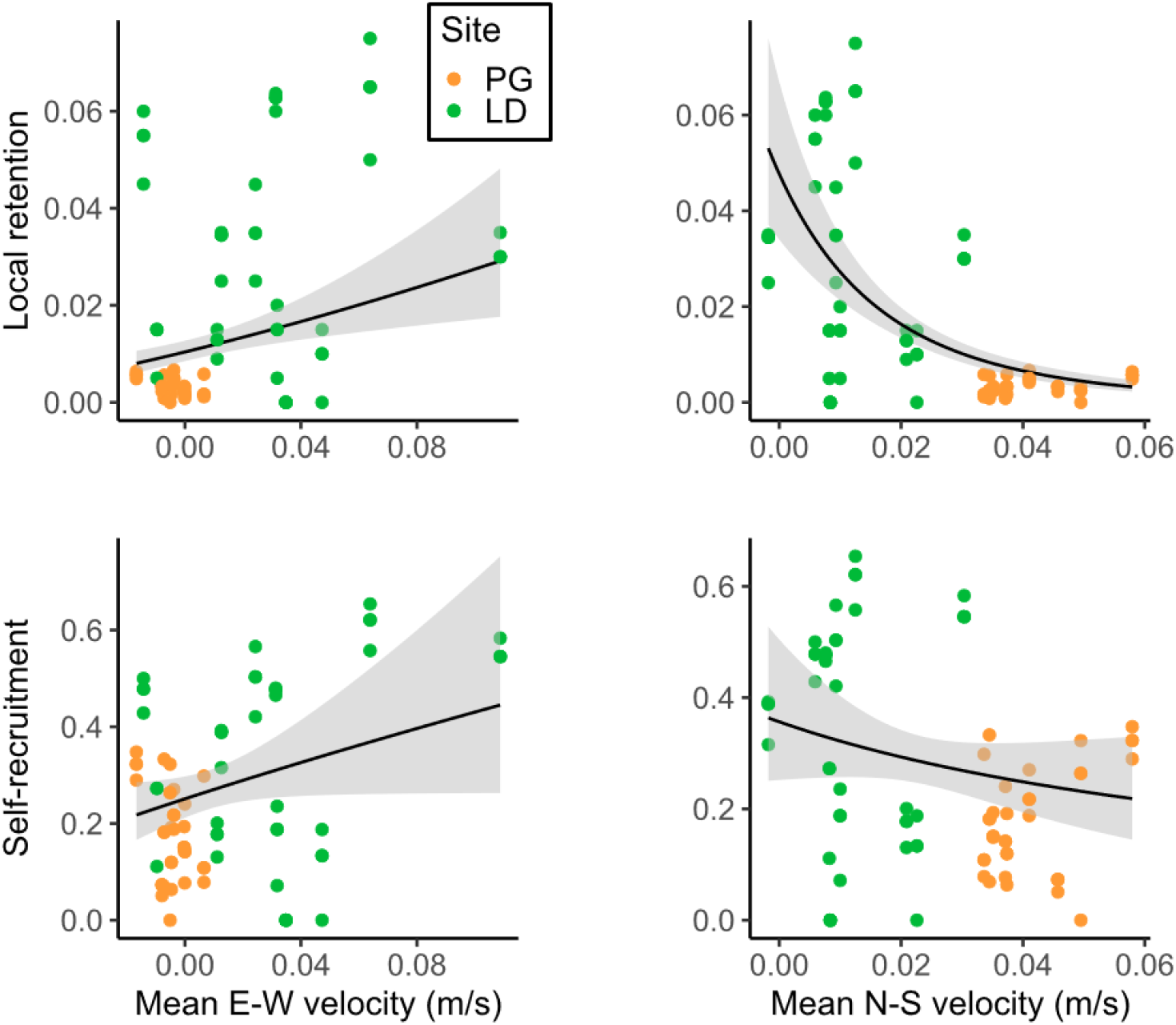
Local retention and self-recruitment in relation to mean E-W and N-S velocities based on larval dispersal and hydrodynamic simulations in Puerto Galera (PG) and Laguindingan (LD). Mean of vertically averaged velocity were calculated over 6 subsequent days from the particle release dates at all grids of the inner domains in PG and LD models. Each value of flow velocity was averaged over all grids of these inner domains.

## Discussion

Using a combined approach of genetic parentage analysis and biophysical modelling, we investigated larval dispersal patterns of two anemonefishes in a semi-enclosed bay of Puerto Galera and open coast of Laguindingan in the Philippines. Although we expected higher self-recruitment in the semi-enclosed setting than in the open coast, results of genetic parentage analysis showed lower self-recruitment in the former than in the latter for two anemonefish species. This was qualitatively consistent with larval dispersal simulations by the biophysical model, which predicted lower local retention and self-recruitment in the semi-enclosed bay than in the open coast. Although our empirical and theoretical estimates of larval dispersal were inferred for about a half year, the results indicate that it is difficult to predict self-recruitment from geographic setting alone and information of hydrodynamics is critical for it.

### Genetic parentage analysis

Many empirical studies have estimated larval dispersal patterns and self-recruitment in coral reef fishes (Catalano et al., 2021; D’Aloia et al., 2013; Planes et al., 2009; Sato et al., 2017), but studies comparing dispersal patterns between different geographic sites are limited. We conducted genetic parentage analysis to compare larval dispersal patterns and self-recruitment between a semi-enclosed bay and open coast in continuous seascapes of the Philippines. Although we expected higher self-recruitment in the semi-enclosed bay of Puerto Galera than in the open coast of Laguindingan, our study obtained an opposite result: self-recruitment in Laguindingan (14–15 %) was higher than that in Puerto Galera (0–2 %) for two anemonefishes. This result was not likely to be due to incomplete sampling of anemonefish in Puerto Galera because we collected genetic samples of more than 95 % breeders there. Our previous research found no effect of patch isolation (mean distance from other anemones) on abundance of the two anemonefishes in the same study area of Puerto Galera (Sato et al. 2014a), which was likely to be due to almost no dispersal connectivity within the study area. Pinsky et al. (2012) predicted self-recruitment is lower in continuous seascapes than patchy seascapes because estimates of self-recruitment can be diluted by high numbers of immigrants from surrounding patches in a continuous seascape, or alternatively, they may be increased by a lack of external recruitment in an isolated patch. Empirical marine dispersal researches have confirmed it by showing higher estimates of self-recruitment on isolated habitats in patchy seascapes such as reefs in island than those on habitats in continuous seascapes (22.6–65.0 % vs 0.2–27.1 %) (Almany et al., 2007; Planes et al., 2009; Saenz-Agudelo et al., 2011; Beldade et al., 2012; Berumen et al., 2012; Catalano et al., 2021). Our estimate of self-recruitment in Puerto Galera was consistent with this prediction, but our estimate (0–2 %) was one of the lowest among previous estimates in continuous seascapes. There are two comparable studies reporting a similar level of self-recruitment for anemonefish species (Nanninga et al., 2015 ; Catalano et al., 2021). The study of Nanninga et al. (2015) was conducted on a small island isolated from any neighboring reefs by an average distance of approximately 10 km in the Red Sea. Authors indicated that their measures of self-recruitment were considerably lower than those of any comparable studies and their result in the patchy seascape was deviated from the norm predicted by Pinsky et al. (2012). They argued that the small size of the study site or selective advantage for emigration from the natal site was a candidate mechanism for their result but admitted these were still pure speculations (Nanninga et al., 2015). Thus, a direct comparison with their study is difficult. Meanwhile, Catalano et al. (2021) reported low self-recruitment (0.23–8.6%) of anemonefish in a continuous seascape along 30 km of coastline in the Philippines, and their minimum value was near to our estimate in Puerto Galera. Although they did not indicate reasons for low self-recruitment in the study site, they found differences in dispersal direction and distance between monsoon seasons. Their results suggested predominantly southwards dispersal in the Northeast Monsoon season (wind blowing out of the northeast) and predominantly northwards dispersal in the Southwest Monsoon season (wind blowing out of the southwest). Our juvenile samples were estimated to disperse as larvae in the Southwest Monsoon season (May to September 2012) in Puerto Galera and in both Northeast (February to April 2013) and Southwest Monsoon (May to July 2013) seasons in Laguindingan. There is one possibility that the larvae released from Puerto Galera in the Southwest Monsoon season were dispersed in a northward direction by wind-driven current, but it is also possible for the larvae released from Laguindingan in this season. Our simulation results showed the velocity magnitude and N-S velocity around Puerto Galera were larger than those off Laguindingan even in the same season (Fig. S5). Therefore, we infer that the general oceanographic features in each site mainly contributed to the contrasting self-recruitment values between them. Such a result was consistent with previous studies that showed that local oceanography was one of the most important determinants of self-recruitment and local retention (Treml et al., 2012).

### Larval dispersal simulations

To examine whether the limited self-recruitment in Puerto Galera relative to Laguindingan is a result of low rates of larval retention or possibly due to an attenuation of self-recruitment estimates via high levels of larval imports from external habitats, we implemented a high-resolution biophysical dispersal model. Our larval dispersal simulations predicted a considerably lower proportion of larvae retained in Puerto Galera (local retention = 0.2–2.1%) than in Laguindingan (0.8–5.3%) under all parameters (Table 1). This lower value of local retention in Puerto Galera can be the main factor for the lower self-recruitment in Puerto Galera (9–47%) than in Laguindingan (22–57%). Multiple regression analysis showed local retention and self-recruitment values decreased with increasing the mean N-S velocity (Fig. 6). Northward currents direct to the offshore in our study sites, and the stronger cross-shore flow may transport anemonefish larvae to the offshore and decrease local retention and self-recruitment values especially in Puerto Galera. Meanwhile, this regression analysis also found local retention and self-recruitment increased with increasing the E-W velocity, which is the along-shore directions in our sites (Fig. 2). Stronger eastward currents around Laguindingan may transport larvae released from Laguindingan to the east, but they are retained within the Macajalar bay and can come back to Laguindingan. Therefore, our analysis may detect the positive relationship between the E-W velocity and local retention or self-recruitment.

We also must consider other possible factors creating the variation in self-recruitment between Puerto Galera and Laguindingan. Larval dispersal simulations found that relative number of recruits from the surrounding locations was higher in Puerto Galera (76.3 recruits from external sources vs 19.6 from internal sources) than in Laguindingan (52.7 vs 31.9) (Table S3), which also led to lower self-recruitment in the former. This patten could be derived if relative number of the external to internal sources was higher in Puerto Galera than in Laguindingan. However, an actual ratio of the external to internal sources was lower in the former ((45 + 34)/5 = 15.8) than in the latter ((15 + 14)/1 = 29.0). Therefore, the ratio of the external to internal sources was not responsible for the variation in self-recruitment between the two sites. Also, a previous study showed that the size of the focal habitat patch had positive effects on self-recruitment (Treml et al., 2012), providing a possibility that variation in sink patch size between Puerto Galera and Laguindingan can affect estimates of self-recruitment. However, our observed patterns were opposite to the prediction of Treml et al. (2012) that there would be higher estimates of self-recruitment in Puerto Galera than in Laguindingan, because the sink patch size was larger in the former. This possibility could be thus excluded, and we conclude that oceanographic feature shapes contrasting patterns of self-recruitment between Puerto Galera and Laguindingan.

Meanwhile, we should not overlook the discrepancy between the simulated and empirical estimates of self-recruitment (original simulation value of 19 % and empirical value of 0–2 % in Puerto Galera; 38 % and 14–15 % in Laguindingan). For both sites, we defined release locations along the shore within a radius of 20 km from the survey area based on the estimated distance at which 90 % of anemonefish larvae recruited (17.6 km) in a previous study, but it also indicated a certain proportion of the larvae dispersed more than 20 km (1–10 %) (Catalano et al., 2021). Therefore, our release locations limited within a radius of 20 km may increase the self-recruitment of larval dispersal simulations. In addition, we set a similar density of particle releases in the study sites and surrounding region based on the assumption of comparable anemonefish density between them. However, there is a possibility of higher anemonefish density in the surrounding regions than in the focal study sites. In that case, larval dispersal simulations get lower values of self-recruitment in both study sites as our simulation showed it in setting the particle releases from the focal locations to a half value (50 particles per station) of the surrounding stations (100 particles per station) (Table 1). In this study, however, we did not measure anemonefish density around the surrounding region. Such additional monitoring will increase the accuracy of prediction of the larval dispersal simulation. Another possibility is that settlement processes that were not included in our model may influence patterns of recruitment. Our dispersal simulation records settlement when an individual gets in contact with a sensory zone after the competency period as in Nanninga et al. (2015). In reality, the larva would still have to actually reach the reef and successfully settle into the benthic habitat. There is strong intra- and interspecific competition of anemonefish, such as prevention of the settlement and eviction of new juveniles by a resident individual, in the settlement process (Buston, 2003; Elliott and Mariscal, 2001; Sato et al., 2014). Juvenile anemonefish are highly vulnerable to predation when they depart from the host (Elliott et al. 1995); this process in the settlement stage can be a major bottleneck or barrier against successful recruitment (Almany and Webster, 2006; Sato et al., 2017). Larval dispersal simulation without such process might overestimate the local retention and self-recruitment.

According to Nanninga et al. (2015), we assessed the influences of different biological parameter inputs on local retention and self-recruitment estimates in both models. As in the previous study, perturbations of individual parameters had variable effects on the estimates of local retention. Although the number of particle releases had a negligible effect in both models and the effect of vertical migration was only apparent in the Laguindingan model, competency period, mortality rate, and sensory zone remarkably affected estimates of local retention in both models. Indeed, the importance and effects of these parameters are consistent with other studies (Cowen et al., 2000, 2006; Nanninga et al., 2015; Treml et al., 2012). We also found effects of competency period, sensory zones, vertical migration, and number of particle releases from the focal areas on the estimates of self-recruitment. The effects of sensory zones and larval mortality had similar trends to a previous result (Treml et al., 2015). However, the importance and effect of the competitive period were somewhat different from our results. Treml et al. (2015) showed that increasing the start of the competitive period decreases the self-recruitment, but an opposite pattern was observed in the results of our Laguindingan model. In addition, passive migration scenario decreased self-recruitment in Laguindingan but increased it in Puerto Galera. Therefore, the effects of these parameters on self-recruitment can be site dependent.

### Implications for management

When considering the design of marine protected areas (MPAs), the size and spacing of MPAs should be determined by considering the spatial distribution of biodiversity and the connectivity of the population via larval dispersal (e.g., Halpernand Warner 2003; Planes et al., 2009). In closed systems, where a higher proportion of larvae are expected to stay in the area and larval imports from outside are limited, a large fraction of their area is designed as an MPA to ensure population persistence (Jossopp and McAllen, 2007). However, present study showed that fish populations in Puerto Galera were not self-sustained and maintained by supply from outside populations. This indicates that even in closed systems, depending on the characteristics of current velocity and direction, protecting several external source areas or extending an MPA to beyond the bay may be required to enhance population persistence. In case of Puerto Galera, implementing MPAs in the western side of Puerto Galera can be a good option because larger number of larvae are predicted to immigrate from that area (Table S3) but few MPAs have been implemented there (Weeks et al., 2010). Or extending the MPA to out of the bay may increase self-persistence of populations within it. On the contrary, a large MPA is more likely to be self-sustaining in open systems as larvae disperse widely. However, in Laguindingan, relatively high self-recruitment was observed despite the small area in the open system, suggesting that fish populations could be maintained by establishing small MPAs even in open coasts where currents are weak.

According to Weeks et al. (2010), which analyzed 985 MPAs in the Philippines, 90% of MPAs in the Philippines have a total area of <1 km^2^. In the Philippines, the different autonomy of each island and sea area, and the narrow range of movement of fishers, make it difficult to set up large MPAs. This is generally the case in many developing countries with coral reefs, especially where communities rely heavily on these reefs for livelihoods (Ban et al., 2011). When considering the design of MPAs, not only geographic shape but also hydrodynamics (current velocity and direction) should be taken into account to determine the placement of small no-take MPAs to ensure the conservation and management of fish populations. Meanwhile, small MPAs are often not effective for highly mobile fish species (Honda et al. 2016). For such fishes, other options, i.e., establishing temporal MPAs on their spawning grounds during a spawning season and/or setting fishing regulations, may be necessary to increase the effectiveness of management.

## Acknowledgements

We are grateful to Yukio Nagahama and Yvette Geroleo of Japan International Cooperation Agency; Darwin I. Baslot, Allyn Duvin S. Pantallano, and Tom G. Genovia of Mindanao State University at Naawan; Kentaro Iwai of Tokyo Institute of Technology for their helpful support in the field survey; and the Municipality of Puerto Galera and staffs of Mindanao State University at Naawan for their cooperation during the field work. We also thank Tanuspong Pokavanich of Kuwait Institute for Scientific Research and Naoki Furuichi of Fisheries Technology Institute for their advice on biophysical modelling, and Hiroyuki Kurokochi of University of the Tokyo for helpful comments on the early version of the manuscript. Collection of samples was permitted as No. 0065-12 and 0073-14 by the Department of Agriculture-Bureau of Fisheries and Aquatic Resources (DA-BFAR) in accordance with Philippine laws and regulations (Republic Act. No. 9147; FAO 233). This research was jointly funded by the Japan Science and Technology Agency (JST), the Japan International Cooperation Agency (JICA) and Science and Technology Research Partnership for Sustainable Development (SATREPS). No conflicts of interest were declared.

