## Supplementary materials for "Hydrodynamics shapes self-recruitment in anemonefishes"

**Supplemental Information** for the article "Hydrodynamics shapes self-recruitment in anemonefishes"

**Genetic analysis for Puerto Galera populations**

*Summary statistics of microsatellite loci*

The individuals were genotyped using 22 microsatellite loci for *A. frenatus* and 24 loci for *A. perideraion* (Table S1). Sato et al. (2014b) developed 12 of the 22 loci for *A. frenatus* and 12 of the 24 loci for *A. perideraion*. Other loci were identified through cross-species amplification of loci developed in previous studies (Beldade et al., 2009; Buston et al., 2007; Liu et al., 2007; Pinsky et al., 2010; Quenouille et al., 2004; Watts et al., 2004). Genomic DNA was extracted, amplified fragments were sequenced and scored according to Sato et al. (2014). Allele frequencies, observed and expected heterozygosity were calculated, and the frequency of null alleles was estimated using CERVUS v. 3.0 (Kalinowski et al., 2007). Deviations from Hardy–Weinberg equilibrium were calculated using GENEPOP v. 4.2 (Rousset, 2008).

The 22 microsatellite markers for *Amphiprion frenatus* and the 24 markers for *A. perideraion* were polymorphic in Puerto Galera populations. Averaged numbers of alleles per locus were 13.2 and 13.3, ranging from 4 to 38 and from 2 to 32, respectively (Table S1). Deviations from the Hardy–Weinberg equilibrium were detected for each seven loci of A. frenatus and A. perideraion ($p < 0.05$). CERVUS analysis estimated relatively high frequencies of the null allele in two loci [F (null) = 0.059 and 0.175] for *A. frenatus*, and four loci [F (null) = 0.075, 0.053, 0.375 and 0.102] for *A. perideraion*. Because the frequency of a null allele over 0.2 biases parentage assignment (Dakin & Avise, 2004), we excluded one locus of *A. perideraion* (Cf8) for the analysis.

Table S1 Characteristics of the 22 and 24 polymorphic microsatellite loci for *Amphiprion frenatus* and *A. perideraion*, respectively. Reported characteristics at each locus are based on all the individuals sampled at Puerto Galera site.

| Locus | N | No. of alleles | $H_O$ | $H_E$ | HWE (P value) | F (null) | References |
|---|---|---|---|---|---|---|---|
| *A. frenatus* (n =206) | | | | | | | |
| Af-01 | 206 | 12 | 0.718 | 0.755 | 0.613 | 0.030 | Sato et al. (2014) |
| Af-02 | 206 | 5 | 0.505 | 0.471 | 0.691 | -0.045 | Sato et al. (2014) |
| Af-03 | 205 | 38 | 0.946 | 0.957 | 0.258 | 0.004 | Sato et al. (2014) |
| AfAp-01 | 205 | 17 | 0.815 | 0.884 | 0.022 | 0.041 | Sato et al. (2014) |
| AfAp-02 | 206 | 23 | 0.816 | 0.919 | <0.001 | 0.059 | Sato et al. (2014) |
| AfAp-03 | 206 | 11 | 0.835 | 0.839 | 0.638 | 0.003 | Sato et al. (2014) |
| AfAp-04 | 203 | 22 | 0.906 | 0.891 | 0.815 | -0.010 | Sato et al. (2014) |
| AfAp-05 | 205 | 12 | 0.766 | 0.797 | 0.109 | 0.022 | Sato et al. (2014) |
| AfAp-07 | 206 | 27 | 0.864 | 0.912 | <0.001 | 0.026 | Sato et al. (2014) |
| AfAp-08 | 205 | 13 | 0.834 | 0.855 | 0.003 | 0.008 | Sato et al. (2014) |
| AfAp-09 | 205 | 6 | 0.600 | 0.566 | 0.830 | -0.039 | Sato et al. (2014) |
| AfAp-10 | 206 | 18 | 0.869 | 0.869 | 0.425 | -0.002 | Sato et al. (2014) |
| 2 | 205 | 4 | 0.098 | 0.103 | 0.356 | 0.021 | Quenouille et al. (2004) |
| 45 | 206 | 18 | 0.874 | 0.868 | 0.335 | -0.006 | Quenouille et al. (2004) |
| 65 | 199 | 4 | 0.698 | 0.669 | 0.791 | -0.023 | Quenouille et al. (2004) |
| LIST-28 | 205 | 15 | 0.663 | 0.694 | 0.009 | 0.022 | Watts et al. (2004) |
| Cf8 | 206 | 5 | 0.510 | 0.502 | 0.396 | -0.014 | Buston et al. (2007) |
| Cf29 | 205 | 18 | 0.883 | 0.892 | 0.029 | 0.003 | Buston et al. (2007) |
| 1578 | 206 | 5 | 0.621 | 0.611 | 0.694 | -0.009 | Liu et al. (2007) |
| A130 | 206 | 10 | 0.150 | 0.208 | 0.002 | 0.175 | Beldade et al. (2009) |
| A115 | 206 | 4 | 0.083 | 0.089 | 0.291 | 0.032 | Beldade et al. (2009) |
| C1 | 206 | 5 | 0.617 | 0.647 | 0.079 | 0.020 | Pinsky et al. (2010) |
| *A. perideraion* (n = 98) | | | | | | | |
| Ap-01 | 98 | 16 | 0.857 | 0.881 | 0.629 | 0.010 | Sato et al. (2014) |
| Ap-02 | 98 | 6 | 0.571 | 0.655 | 0.071 | 0.075 | Sato et al. (2014) |
| AfAp-01 | 98 | 14 | 0.776 | 0.805 | 0.035 | 0.013 | Sato et al. (2014) |

| | | | | | | | |
|---|---|---|---|---|---|---|---|
| AfAp-02 | 98 | 16 | 0.735 | 0.828 | 0.297 | 0.053 | Sato et al. (2014) |
| AfAp-03 | 98 | 11 | 0.786 | 0.833 | 0.313 | 0.026 | Sato et al. (2014) |
| AfAp-04 | 98 | 13 | 0.908 | 0.873 | 0.433 | -0.023 | Sato et al. (2014) |
| AfAp-05 | 98 | 13 | 0.878 | 0.864 | 0.059 | -0.012 | Sato et al. (2014) |
| AfAp-06 | 98 | 6 | 0.520 | 0.561 | 0.160 | 0.030 | Sato et al. (2014) |
| AfAp-07 | 98 | 16 | 0.898 | 0.873 | 0.056 | -0.021 | Sato et al. (2014) |
| AfAp-08 | 98 | 9 | 0.867 | 0.830 | 0.542 | -0.029 | Sato et al. (2014) |
| AfAp-09 | 98 | 4 | 0.418 | 0.423 | 0.864 | 0.003 | Sato et al. (2014) |
| AfAp-10 | 98 | 17 | 0.908 | 0.882 | 0.234 | -0.020 | Sato et al. (2014) |
| 2 | 98 | 3 | 0.347 | 0.331 | 0.845 | -0.031 | Quenouille et al. (2004) |
| 45 | 97 | 32 | 0.938 | 0.944 | 0.090 | 0.001 | Quenouille et al. (2004) |
| Cf8 | 98 | 5 | 0.020 | 0.041 | <0.001 | 0.375 | Buston et al. (2007) |
| Cf29 | 96 | 18 | 0.854 | 0.857 | 0.515 | -0.004 | Buston et al. (2007) |
| 915 | 98 | 9 | 0.643 | 0.633 | 0.352 | -0.006 | Liu et al. (2007) |
| 1578 | 98 | 5 | 0.367 | 0.359 | 0.023 | -0.031 | Liu et al. (2007) |
| A115 | 98 | 24 | 0.714 | 0.765 | <0.001 | 0.043 | Beldade et al. (2009) |
| A130 | 98 | 21 | 0.847 | 0.883 | 0.390 | 0.016 | Beldade et al. (2009) |
| D1 | 98 | 13 | 0.878 | 0.839 | 0.450 | -0.024 | Beldade et al. (2009) |
| D103 | 97 | 23 | 0.732 | 0.901 | <0.001 | 0.102 | Beldade et al. (2009) |
| B6 | 98 | 14 | 0.867 | 0.871 | 0.058 | 0.000 | Pinsky et al. (2010) |
| C1 | 93 | 2 | 0.065 | 0.063 | 1.000 | -0.008 | Pinsky et al. (2010) |

$H_O$ observed heterozygosity, $H_E$ expected heterozygosity, HWE deviation from Hardy–Weinberg equilibrium test, F (null) frequency of null alleles.

*Simulation for parentage analysis*

To assess the information sufficiency of our markers for the accurate reconstruction of parental assignment, we used the simulation module in COLONY (Wang, 2013). The module simulated juvenile genotypes with a predefined parentage and sibship structure, based on a given marker number, allele frequencies, and an assumed mating matrix. It then returned a metric of the accuracy of parentage assignments (Muralidhar et al., 2014). We used parameters identical to the original COLONY run to simulate juveniles at the study site and to determine the confidence in our parentage assignments.

The simulation results indicated a 0 % of a Type I error (probability of assigning to a false parent) as well as a 0 % chance of a Type II error (probability of falsely excluding a parent when it was in the sample) for both species.

### Hydrodynamic model

#### *Hydrodynamic model configuration*

Using Delft3D-Flow, a 3D-capable hydrodynamic simulation program (Deltares, 2014a), we developed two hydrodynamic models, i.e., the Puerto Galera model extending from 13.38 to 13.93 N and from 120.62 to 121.26 E, and the Laguindingan model extending from 8.41 to 8.99 N and from 124.07 to 124.78 E (Fig. 2). Both models were comprised of two computational domains using one-way nesting technique to properly incorporate the effects of the complicated topographic setup and tidal fluctuation. For the Puerto Galera model, the Verde Island Passage area was an outer domain with a cell size of 900 m and the Puerto Galera domain with a cell size of 100 m was the inner domain. For the Laguindingan model, the Bohol Sea area was an outer domain with a cell size of 900 m, and the Laguindingan area with a cell size of 100 m was the inner domain. Thirteen and fifteen vertical Z-layers were used with increasing the thickness from the surface down to the bottom for Puerto Galera and Laguindingan models, respectively. The General Bathymetric Chart of the Oceans (GEBCO: http://www.gebco.net/) data were supplemented with bathymetry data of Pokavanich (2009) and echo sounder measurements in both areas, where coral reefs make topography highly variable. The simulated water levels from the PH model (Pokavanich 2009) were recorded in five-minute intervals, bilinearly interpolated, and applied along the open boundaries of the Verde Island Passage and Bohol Sea domains. The daily average reanalysis data of salinity and temperature from JCOPE2 (Miyazawa et al., 2009) and Hybrid Coordinate Modeling System analysis data (HYCOM GLBu0.08) (https://hycom. org/dataserver/glb-analysis) were used for the initial and boundary conditions of Verde Island Passage and Bohol Sea domains, respectively. The domain decomposition function, which can run simulations of separate domains in parallel, was used to smoothly communicate the boundary condition between two computational domains of both models (Deltares, 2014a). The water surface was forced with a spatially uniform wind, whose data were derived from a weather station (HOBO, Onset Computer Corporation, USA) at P1 in Puerto Galera and L4 in Laguindingan (Fig. 2) and from a climatological database (http://www.windfinder.com) for Calapan of Mindoro Island, 25 km southeast from the Puerto Galera, and those for Cagayan de Oro Airport of Mindanao Island, 27 km southeast from the Laguindingan. We used the wind data at the several points because data at one point could not cover the whole study period. To calculate the heat exchange through the surface, an ocean heat flux model was used to prescribe the relative humidity, air temperature, and fraction of cloud coverage (Deltares, 2014a). The relative humidity and air temperature were recorded by the weather station, while the fraction of cloud coverage was obtained from the NASA earth observations (http://neo.sci.gsfc.nasa.gov/). Because evaporation and precipitation cannot be neglected in tropical regions (Deltares, 2014a), precipitation data and calculated evaporation were combined with the ocean heat flux model. For the ocean heat flux model, Secchi depth was set to 12 m for the Puerto Galera model and 16 m for the Laguindingan model according to the Reef Check monitoring data. The Dalton number for the evaporative heat and Stanton number for the heat convection were also prescribed to 0.001 and 0.002, respectively, for both models following the ranges as in Li and Reidenbach (2014). The background horizontal eddy viscosity and diffusivity on Verde Island Passage and BH domains were set to be spatially variable, whereas those on the Puerto Galera domain were set to 1 and 10 $m^2 s^{-1}$ and those on Laguindingan domain were set to 5 and 10 $m^2 s^{-1}$. In addition to the background values, the viscosity and diffusivity were calculated using the Horizontal Large Eddy Simulation (Deltares, 2014a). The background vertical eddy viscosity and diffusivity were set to $10^{-4}$ $m^2 s^{-1}$ for both models (Miyake et al., 2011). The additional viscosity and diffusivity were computed using the $k$-$\varepsilon$ turbulence model.

*Model validation*

Model performance was evaluated by comparing simulation results with in-situ measurements at each study site (P1 and P2 in Puerto Galera and L1–L4 in Laguindingan) for water level and flow velocity. We calculated the model skill score as defined by Willmott (1981) for every time period that had both modelled and measured values available. The perfect agreement of observed and modeled data yields a model skill score of one, whereas complete disagreement yields the score of zero.

Fig. S1 and S2 showed simulation results and field measurements for water level, East-West, and North-South components of flow velocity in Puerto Galera and Laguindingan. The modeled water level agreed well with observed values, the skill scoring more than 0.94 at the stations in the two sites (Table S2). The velocity plots showed a similar trend between the modeled and observed velocity in most stations in Puerto Galera and Laguindingan (Figs. S1, S2), but some discrepancy in amplitude or timing were observed. However, the skill score of 0.5 and greater, at half and majority stations in Puerto Galera and Laguindingan, respectively (Table S2), indicated each hydrodynamic model well reproduced oceanographic feature around each site.

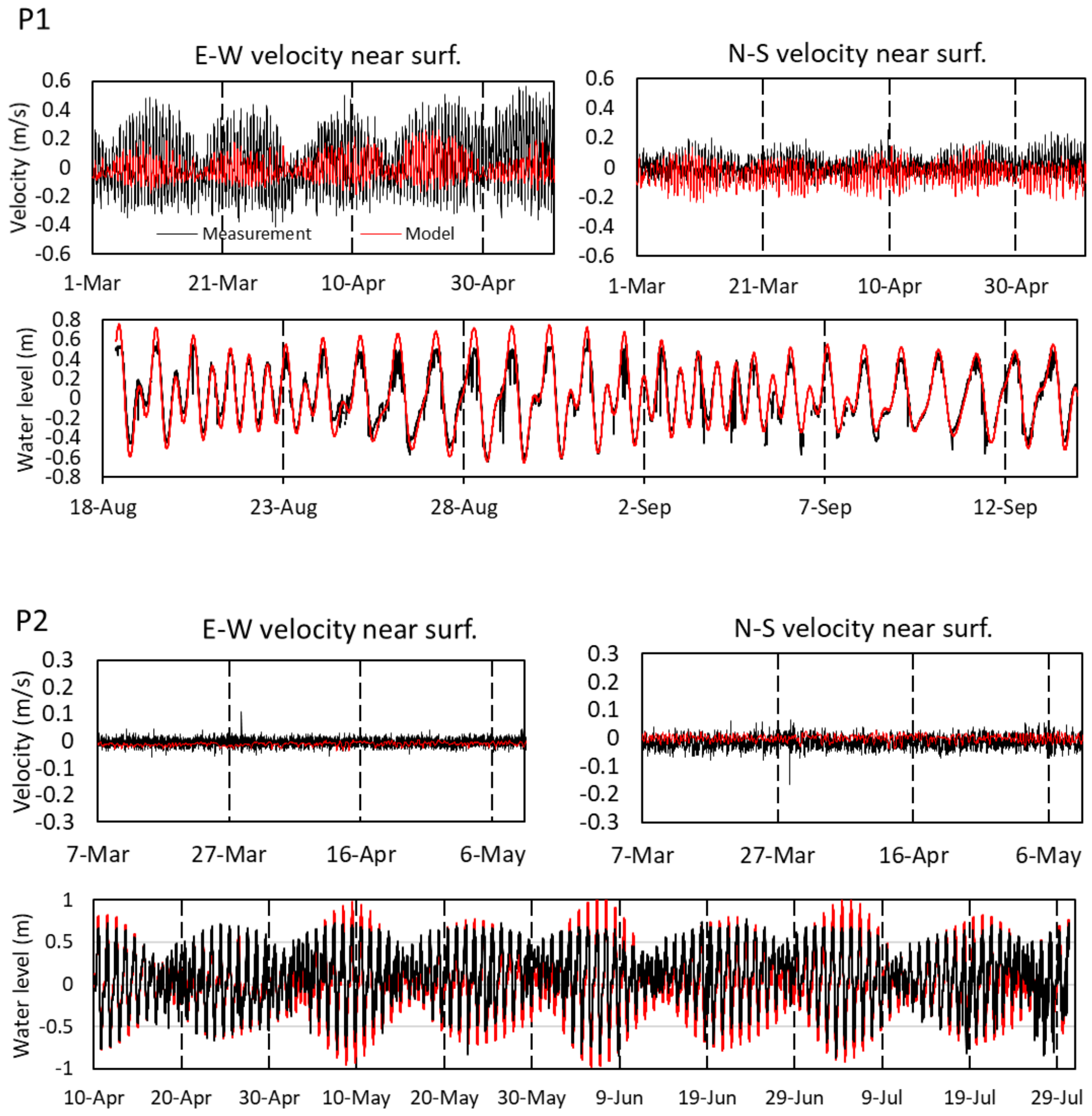


Fig. S1 Comparisons of time-series of measured and modeled near-surface flow velocity and water level at P1 and P2 in Puerto Galera. Near-surface flow velocity was measured by ADCPs, while water level was recorded by water level loggers.

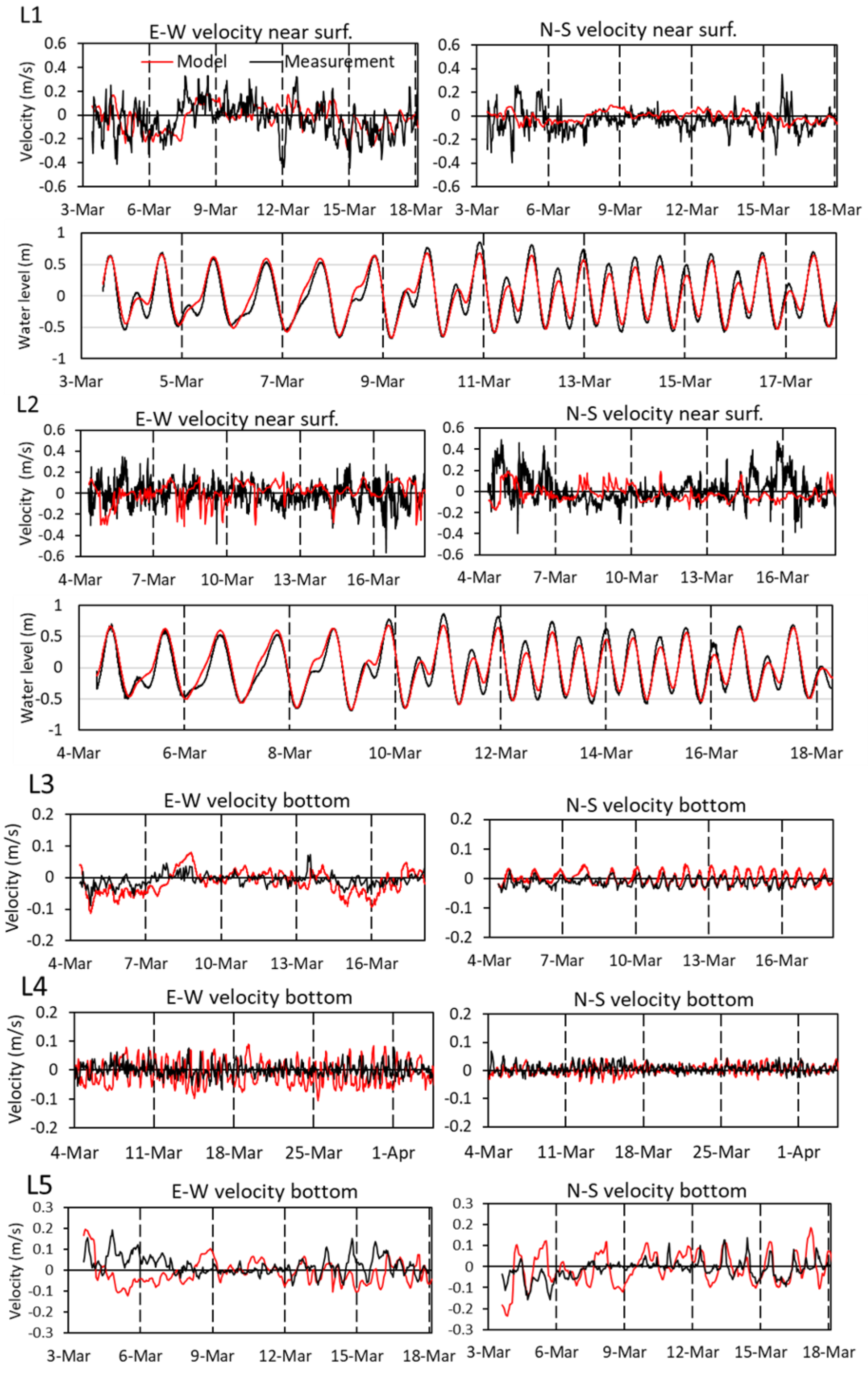


Fig. S2 Comparisons of time-series of measured and modeled near-surface flow velocity and water level at L1 and L2; near-bottom flow velocity at L3, L4, and L5 in Laguindingan.

Table S2. Wilmott agreement scores (Wilmott 1981) for the Puerto Galera (PG) and Laguidingan (LD) models at selected points.

| Model | Station | Instrument | Layer | Period | EW-component velocity | NS-component velocity | Water level |
|---|---|---|---|---|---|---|---|
| PG model | P1 | ADCP/Water level logger | Near surface | Feb 29–Jul 4, 2012/Aug 18–Sep 15*, 2012 | 0.66 | 0.55 | 0.968 |
| | P2 | ADCP/Water level logger | Near surface | Mar 7–Jul 31, 2012/Apr 10–Jul 30, 2012 | 0.36 | 0.37 | 0.942 |
| LD model | L1 | ADCP/Water level logger | Near surface | Mar 4–18, 2013 | 0.58 | 0.28 | 0.978 |
| | L2 | ADCP/Water level logger | Near surface | Mar 4–18, 2013 | 0.28 | 0.57 | 0.978 |
| | L3 | Compact-EM | Bottom | Mar 4–18, 2013 | 0.62 | 0.79 | - |
| | L4 | Infinity-EM | Bottom | Mar 3–Apr 4, 2013 | 0.34 | 0.57 | - |
| | L5 | Infinity-EM | Bottom | Mar 3–18, 2013 | 0.51 | 0.63 | - |

*Water level at P1 is only available from August to September 2012.

*Modelling results*

Fig. S3 and S4 shows residual velocity of vertically averaged flow for the periods from the particle releases (day 0) to start of settling (day 6) around Puerto Galera and Laguindingan. These values were extracted from hydrodynamic simulations of each model. During all the periods, strong northward and westward flows were observed in the eastern side and offshore of Puerto Galera, respectively, whereas flow speed was gradually reduced to the shallower waters of the inner domain (Fig. S3). On the contrary, strong eastward flows were observed in the offshore of Laguindingan for the periods of May 25–31 and Jun 9–15, and relatively weak flows were found there for the remaining periods (Fig. S4). Flow speed was reduced to the shallower waters of the inner domain as in the Puerto Galera model.

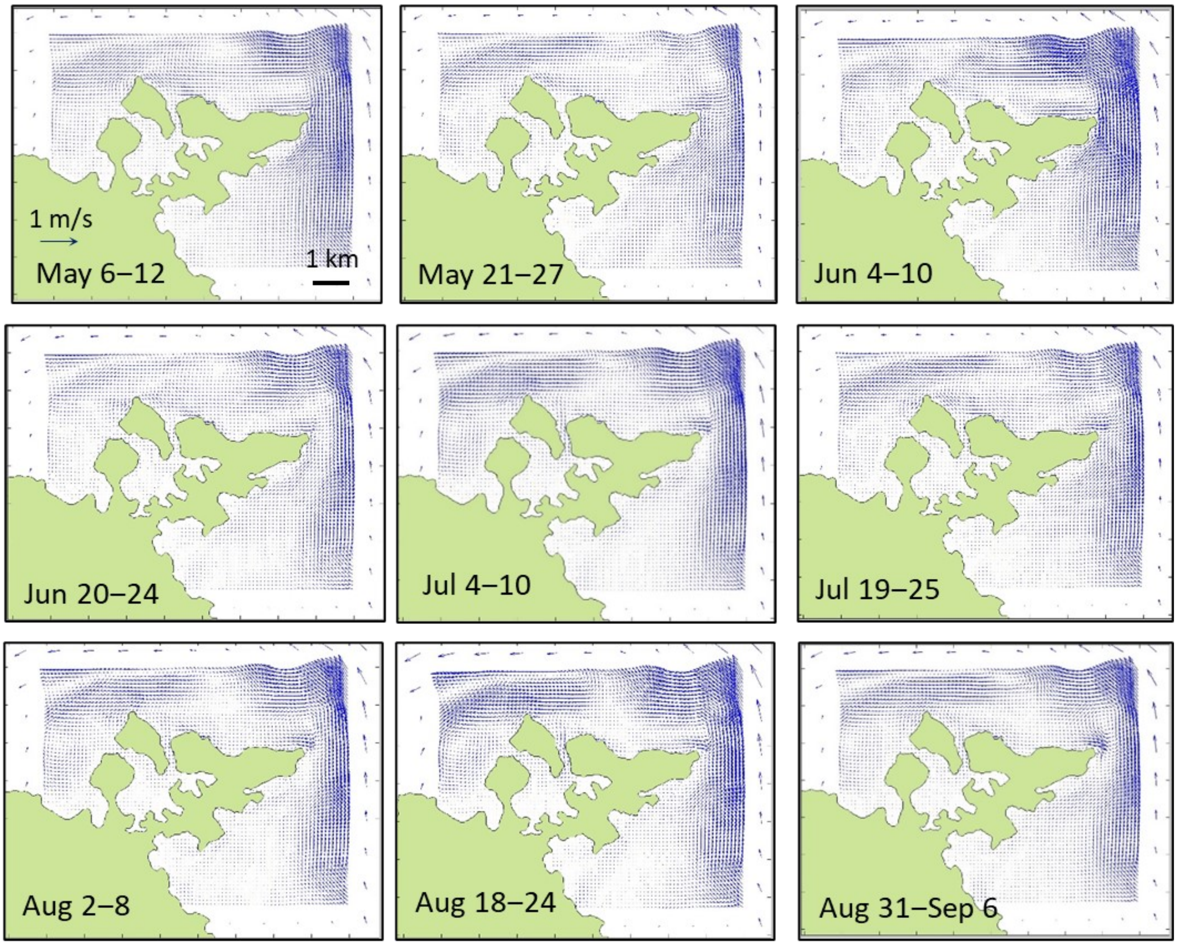


Fig. S3 Residual velocity of vertically averaged flow around Puerto Galera. Arrows indicate the magnitude and directions of vertically averaged flow.

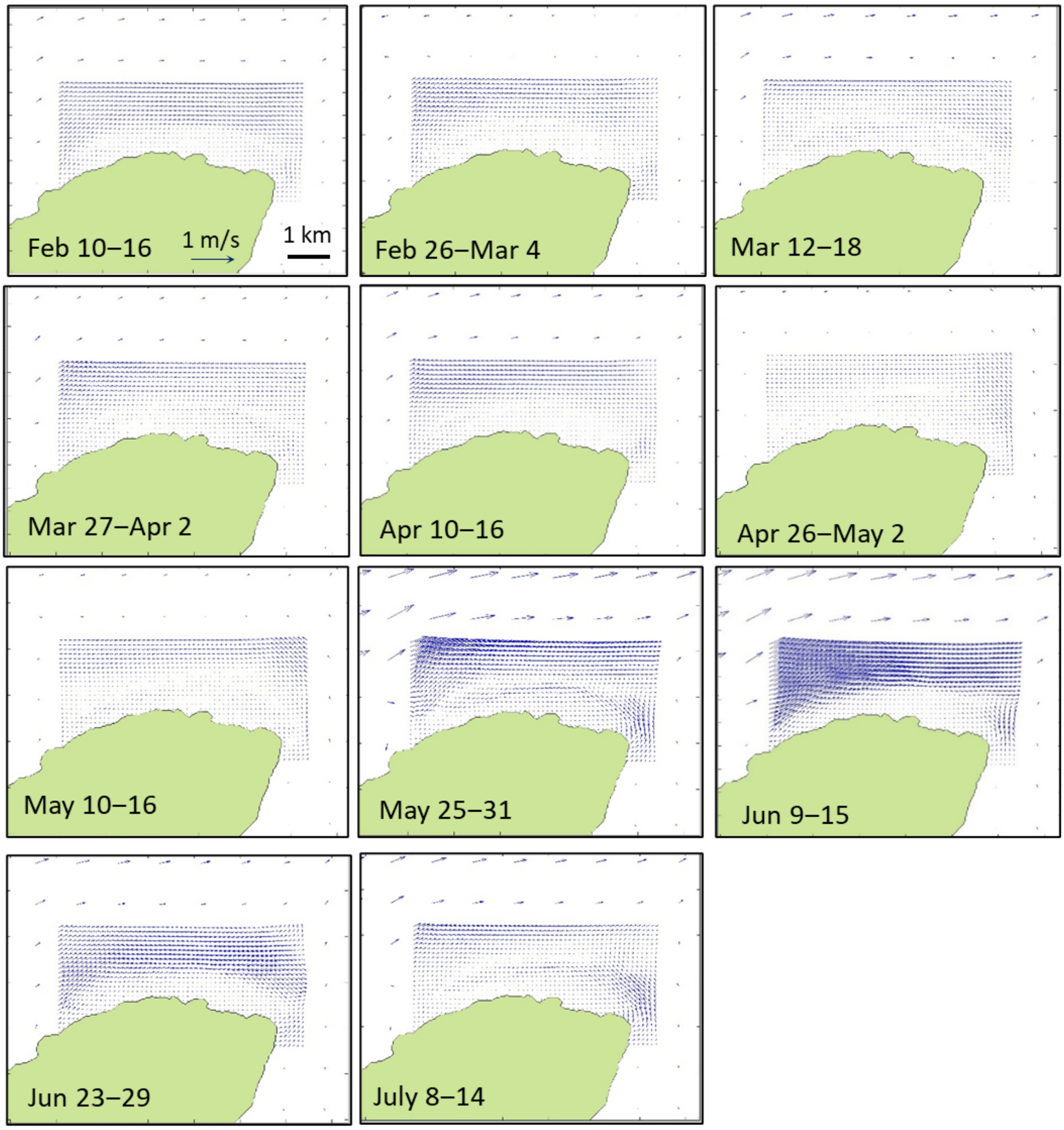


Fig. S4 Residual velocity of vertically averaged flow around Laguindingan. Arrows indicate the magnitude and directions of vertically averaged flow.

**Larval dispersal simulation**

We performed larval dispersal simulations for anemonefish juveniles (<30 mm) at the two sites using Delft3D-PART (Deltares, 2014b). The horizontal dispersion coefficient was set to the same value as the background horizontal dispersal coefficient in the FLOW (Miyake et al., 2011). The vertical dispersion coefficient was calculated using the "depth-averaged algebraic" method, which assumed that the vertical dispersion is constant over depth but varies with location due to spatial variation in depth (Deltares, 2014b). Time step for the particle-tracking calculation was set to 30 minutes. PART simulated larval dispersal from the study site and external surrounding sources into the focal area of the study site in each model domain.

According to Ross (1978), hatching peaks of *Amphiprion* occurred at 1.5 hours after sunset during new and full moons in the tropical region, thus each particle was released between 19:30 and 24:00 on the day of spring tide in the models. Collected juveniles of anemonefishes (<30 mm) in both sites were expected to hatch within four months (Ochi, 1986). In Puerto Galera, genetic samples of the anemonefishes were collected in September 2012, therefore the particle releases in the model were started four months prior to the sample collection. Thus, we set dates of particle releases on May 6, May 21, June 4, June 20, July 4, July 19, August 2, August 18, and August 31, 2012. In Laguindingan, the anemonefish was collected from the end of May to July, 2013 (Sato et al., 2017), and the dates were set on February 10, February 26, March 12, March 27, April 10, April 26, May 10, May 25, June 9, June 23, and July 8, 2013. Based on the distance at which 90 % of anemonefish larvae recruit (17.6 km) as estimated by Catalano et al. (2021), we utilized the 20 km radius to set source sites around each study site. In the Puerto Galera model, a total of 75,600 particles were released from 84 locations (100 particles × 9 spring tides × 84 locations) along the shore within a radius of 20 km from the anemonefish survey area in Puerto Galera (Fig. 4). Meanwhile, in the Laguindingan model, we released 33,000 particles from 30 locations (100 particles × 11 spring tides ×

30 locations) along the shore within a radius of 20 km from the survey area of Laguindingan (Fig. 5). The release locations around the focal study sites of both models were arranged at a regular interval with a density of 100 releases/1800 m per spring tide.

We assessed the model's sensitivity to different biological parameter inputs according to Nanninga et al. (2015). Five parameters were examined for their effect on local retention and self-recruitment: competitive period [6 days (±2)], horizontal sensory zone [1 (+1/–0.5) km radius], larval mortality [half-life of 6 (±2) days], number of particle releases from the study site [100 particle releases (+100 particles/–50 particles) per source location], and vertical migration behavior (passive and empirical). The empirical migration behavior was based on distribution patterns measured via plankton surveys in the same family (Huebert et al., 2011; Irisson et al., 2010). This empirical migration of larvae includes diel and ontogenetic vertical migration, which was simulated using a function of settling velocity ($V_t$) in the PART based on the following equation (Dickey-Collas et al., 2009):

$$V_t = A_0 + A_1 \sin(\frac{2\pi(t-6)}{24})$$

The settling velocity of larvae follows a sine curve variation in time with an amplitude ($A_1$) of $5.0 \times 10^{-4}$ m/s, simulating diel vertical movement as larvae move downward and upward to the water surface during the day and night, respectively (Huebert et al., 2011). Ontogenetic depth shift from deep to shallow water during the larval stage of Pomacentridae (Irisson et al., 2010) was also simulated by setting constant settling velocity ($A_0$) at $1.0 \times 10^{-3}$ m/s from days 0 to 6 and $5.0 \times 10^{-4}$ m/s from days 7 to 15. Here, the positive value of the velocity ($V_t$) indicates the settling speed of larvae while the negative value means the ascent speed.

*Comparisons of local retention, self-recruitment, and hydrodynamic characteristics between Puerto Galera and Laguindingan*

To compare dispersal patterns and hydrodynamic characteristic between Puerto Galera and Laguindingan, we made boxplots of the local retention, self-recruitment, velocity magnitude, East–West (E–W), and North–South (N–S) velocities of modeling results in each site in the same season (from May to July). We used the local retention and self-recruitment values of the original scenario for particle releases in May, June, and July [n =18 (6 spring tides × 3 replicates) for Puerto Galera and n =15 (5 spring tides × 3 replicates) for Laguindingan]. The velocity magnitude, as well as the E–W and N–S velocities were averaged over 6 subsequent days from each date of particle releases in the same season at all the grids of each inner domain. Then, these values were averaged over these grids and used for making the plots [n = 6 (6 spring tides) for Puerto Galera and n = 5 (5 spring tides) for Laguindingan].

Boxplots of Fig. S5 showed that the local retention and self-recruitment values in May, June, and July were also higher in Laguindingan than in Puerto Galera. The magnitude and N-S component of flow velocity was higher in Puerto Galera than in Laguindingan, whereas the mean E–W velocity was higher in Laguindingan.

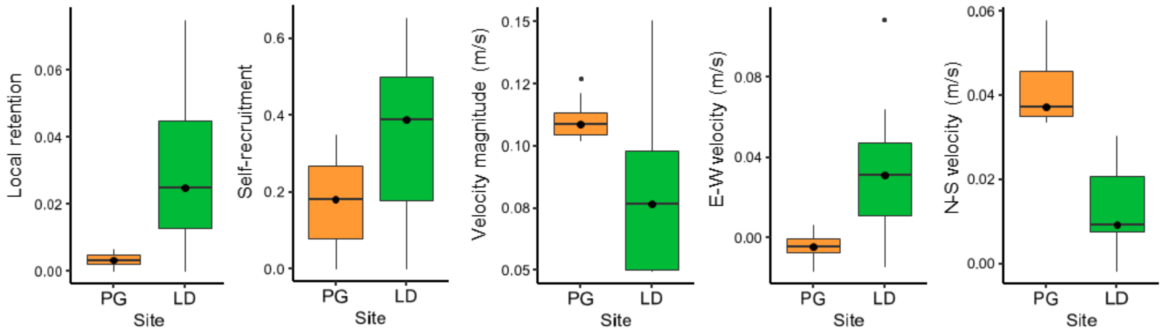


Fig. S5 Boxplots comparing local retention, self-recruitment, velocity magnitude, East–West (E–W), and North–South (N–S) velocities between Puerto Galera (PG) and Laguindingan (LD) in May, June, and July. These values were obtained from larval dispersal and hydrodynamic simulations of Delft3D-Flow and-Part.

Table S3. Numbers of sources, released and settled particles, transport success (%), and self-recruitment in each study area of Puerto Galera (PG) and Laguindingan (LD), predicted in the original scenario of larval dispersal simulations using Delft3D PART.

| Source region | No. of sources | No. of released particles | No. of settled particles in the study area | Transport success (%)* | Self-recruitment |
|---|---|---|---|---|---|
| PG model | | | | | |
| PG | 5 | 4500 | 18.0 | 0.40 | 18.0/(18.0+4.4+71.9)=0.19 |
| East | 45 | 40500 | 4.4 | 0.01 | |
| West | 34 | 30600 | 71.9 | 0.24 | |
| LD model | | | | | |
| LD | 1 | 1100 | 32.3 | 2.93 | 32.3/(32.3+48.6+4.1)=0.38 |
| East | 15 | 16500 | 48.6 | 0.32 | |
| West | 14 | 15400 | 4.1 | 0.03 | |

*Proportion of total particles from each source region that reached the study area. Transport success was calculated as number of settled particles divided by the total released particles from each source region. This value at Puerto Galera and Laguindingan is similar to local retention of each site.
